# SARS-CoV-2 infects human pluripotent stem cell-derived cardiomyocytes, impairing electrical and mechanical function

**DOI:** 10.1101/2020.08.30.274464

**Authors:** Silvia Marchiano, Tien-Ying Hsiang, Ty Higashi, Akshita Khanna, Hans Reinecke, Xiulan Yang, Lil Pabon, Nathan J. Sniadecki, Alessandro Bertero, Michael Gale, Charles E. Murry

**Author notes:** Corresponding authors, contacts: Charles E. Murry, Michael Gale Jr.

## Abstract

Global health has been threatened by the COVID-19 pandemic, caused by the novel severe acute respiratory syndrome coronavirus (SARS-CoV-2)^1^. Although considered primarily a respiratory infection, many COVID-19 patients also suffer severe cardiovascular disease^2–4^. Improving patient care critically relies on understanding if cardiovascular pathology is caused directly by viral infection of cardiac cells or indirectly *via* systemic inflammation and/or coagulation abnormalities^3,5–9^. Here we examine the cardiac tropism of SARS-CoV-2 using human pluripotent stem cell-derived cardiomyocytes (hPSC-CMs) and three-dimensional engineered heart tissues (3D-EHTs). We observe that hPSC-CMs express the viral receptor ACE2 and other viral processing factors, and that SARS-CoV-2 readily infects and replicates within hPSC-CMs, resulting in rapid cell death. Moreover, infected hPSC-CMs show a progressive impairment in both electrophysiological and contractile properties. Thus, COVID-19-related cardiac symptoms likely result from a direct cardiotoxic effect of SARS-CoV-2. Long-term cardiac complications might be possible sequelae in patients who recover from this illness.

## Main text

With over 20 million people affected worldwide, the outbreak of SARS-CoV-2 has already left its permanent mark on human history^1,10^. Cardiovascular complications, including worsening of preexisting conditions and onset of new disorders, significantly contribute to the increasing mortality of COVID-19 patients, but the underlying mechanisms of cardiopathology are unclear^2,11–13^. Upon lung infection, the uncontrolled release of inflammatory cytokines, termed “cytokine storm”, could induce multi-organ damage, ultimately leading to organ failure and worsening of pre-existing cardiovascular disorders^5,6,8,14^. Moreover, COVID-19 is associated with coagulopathies, which also can induce heart damage^3,5,7^. Lastly, SARS-CoV-2 could directly mediate heart injury by entering cardiomyocytes *via* binding of the viral spike glycoprotein to its extracellular receptor, angiotensin I converting enzyme 2 (ACE2)^9,15,16^. An increasing number of reports have showed presence of SARS-CoV-2 genome in the heart and signs of viral myocarditis in COVID-19 infected individuals, including asymptomatic cases, indicating that SARS-CoV-2 could exhibit cardiac tropism and thus directly impair cardiac function^17–20^. Myocardial infarction, arrhythmias, and heart failure are the most common cardiovascular complications observed in COVID-19 patients^12,21,22^. In this study we used human pluripotent stem cell-derived cardiomyocytes (hPSC-CMs) and engineered human heart tissues as platforms to study SARS-CoV-2 cardiac biology^23,24^, to clarify the functional changes behind these COVID-19-related cardiovascular symptoms.

The susceptibility to SARS-CoV-2 infection depends on the expression of the viral receptor ACE2^9,15,16^. We found that *ACE2* is transcriptionally upregulated during cardiac differentiation of both RUES2 embryonic stem cells-derived cardiomyocytes (hESC-CMs; Fig. 1a) and WTC11c induced pluripotent stem cells-derived cardiomyocytes (hiPSC-CMs; Supplementary Fig. 1a). Single-cell RNA-sequencing analysis detected *ACE2* mRNA in ~9% of hESC-CMs, indicating low and/or transitory transcriptional activation (Fig. 1b). A larger fraction of cells expressed moderate to high levels of endosomal cysteine proteases *CTSB* (cathepsin B; ~71.0%) and *CTSL* (cathepsin L; ~46.0%). Detection of these factors is relevant because they can cleave the spike glycoprotein and lead to endomembrane fusion-mediated release of SARS-CoV-2 genome inside the cytoplasm^15,25,26^. Importantly, these viral processing factors were often co-expressed with *ACE2* (Supplementary Fig. 1b). Although viral entry also can be mediated by *TMPRSS2*^15^, this transmembrane serine protease was not detectable in hESC-CMs (Supplementary Fig. 1c), as also reported for the adult human heart^27^. Thus, the ACE2-endosomal viral entry pathway may be the only route for virus entry in cardiomyocytes. Indeed, *PIKFYVE*, another endosomal viral processing factor, was also broadly expressed in hESC-CMs (Supplementary Fig. 1c)^26^. Despite the low levels of mRNA, ACE2 protein was strongly expressed in hPSC-CMs derived from multiple lines (RUES2 female hESCs, H7 female hESCs, and WTC11c male hiPSCs), reaching levels comparable to those of VERO cells, a primate kidney epithelial line with established SARS-CoV-2 tropism (Fig. 1c)^15^. Collectively, we concluded that hPSC-CMs express proteins that render them susceptible to SARS-CoV-2 infection^23,24^.

**Figure 1.**
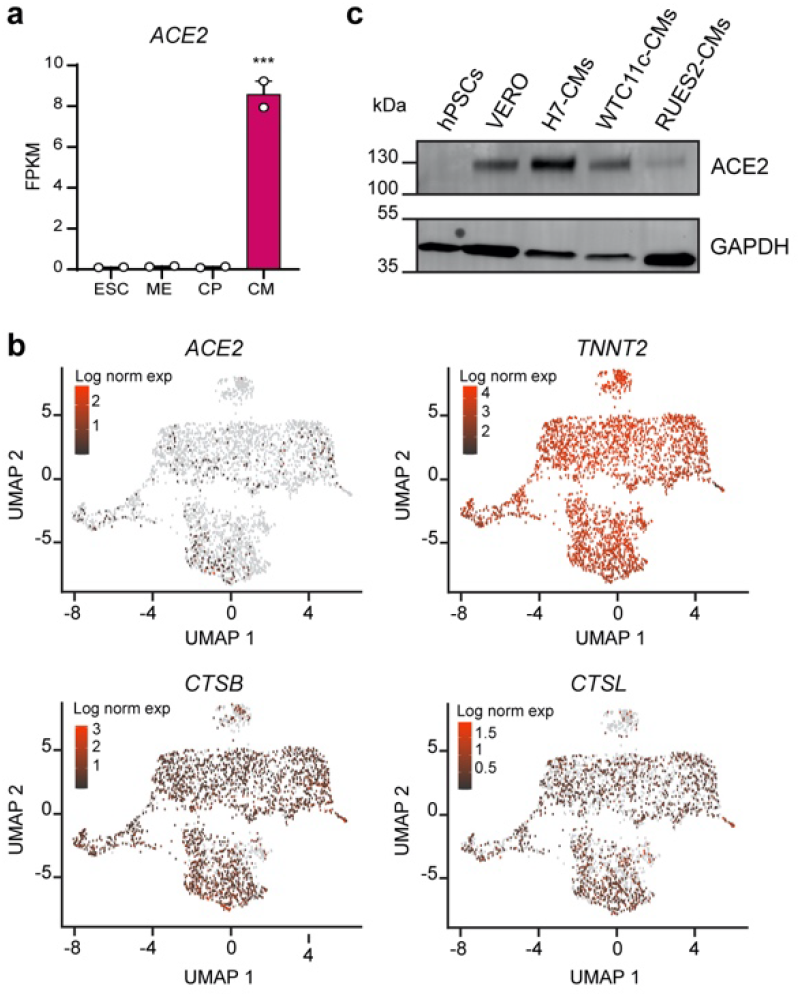
hPSC-CMs express SARS-CoV-2 receptors and processing factors. (**a**) RNA-seq during RUES2 hESC-CM differentiation. *ACE2* is quantified as fragments per kilobase of transcript per million mapped reads (FPKM). Mean ± SEM of two independent experiments. ESC: embryonic stem cells (day 0); ME: mesoderm (day 2); CP: cardiac progenitors (day 5); CM: cardiomyocytes (day 14). Differences *versus* ESC by one-way ANOVA followed by Sidak correction for multiple comparisons (*** = p < 0.001). (**b**) sc-RNA-seq gene expression heatmaps from RUES2 hESC-CMs after dimensionality reduction through Uniform Manifold Approximation and Projection (UMAP). *TNNT2* provides a pan-cardiomyocyte marker. (**c**) Western blot for ACE2 in hPSC-CMs from multiple lines. hPSCs: negative control; VERO cells: positive control.

Since H7 and WTC11c-derived hPSC-CMs showed the highest levels of ACE2, we tested their functional susceptibility to SARS-CoV-2. For this, we incubated highly-pure hPSC-CMs (> 80%; Supplementary Fig. 2a) with SARS-CoV-2/Wa-1 strain at multiplicity of infection (MOI; i.e. the number of infectious viral particles per cell) of either 0.1 (requiring propagation of the virus within the cells and secondary infection of others) or 5 (aiming to infect all the cells at the same time). We observed marked and disseminated viral cytopathic effects in both H7 hESC-CMs and WTC11c hiPSC-CMs. These effects were accelerated at 5 MOI, as expected (Fig. 2a and Supplementary Fig. 2b). Most notably, cessation of beating and cell death started as early as at 48 hours post infection (HPI) in both cell lines, with more pronounced effects for H7 cardiomyocytes. Immunofluorescence staining of SARS-CoV-2 nucleocapsid protein revealed substantial presence of viral factors in the cytoplasm of both H7 hESC-CMs and WTC11c hiPSC-CMs (Fig. 2b and Supplementary Fig. 2c) confirming that hPSC-CMs can be infected directly by SARS-CoV-2.

**Figure 2.**
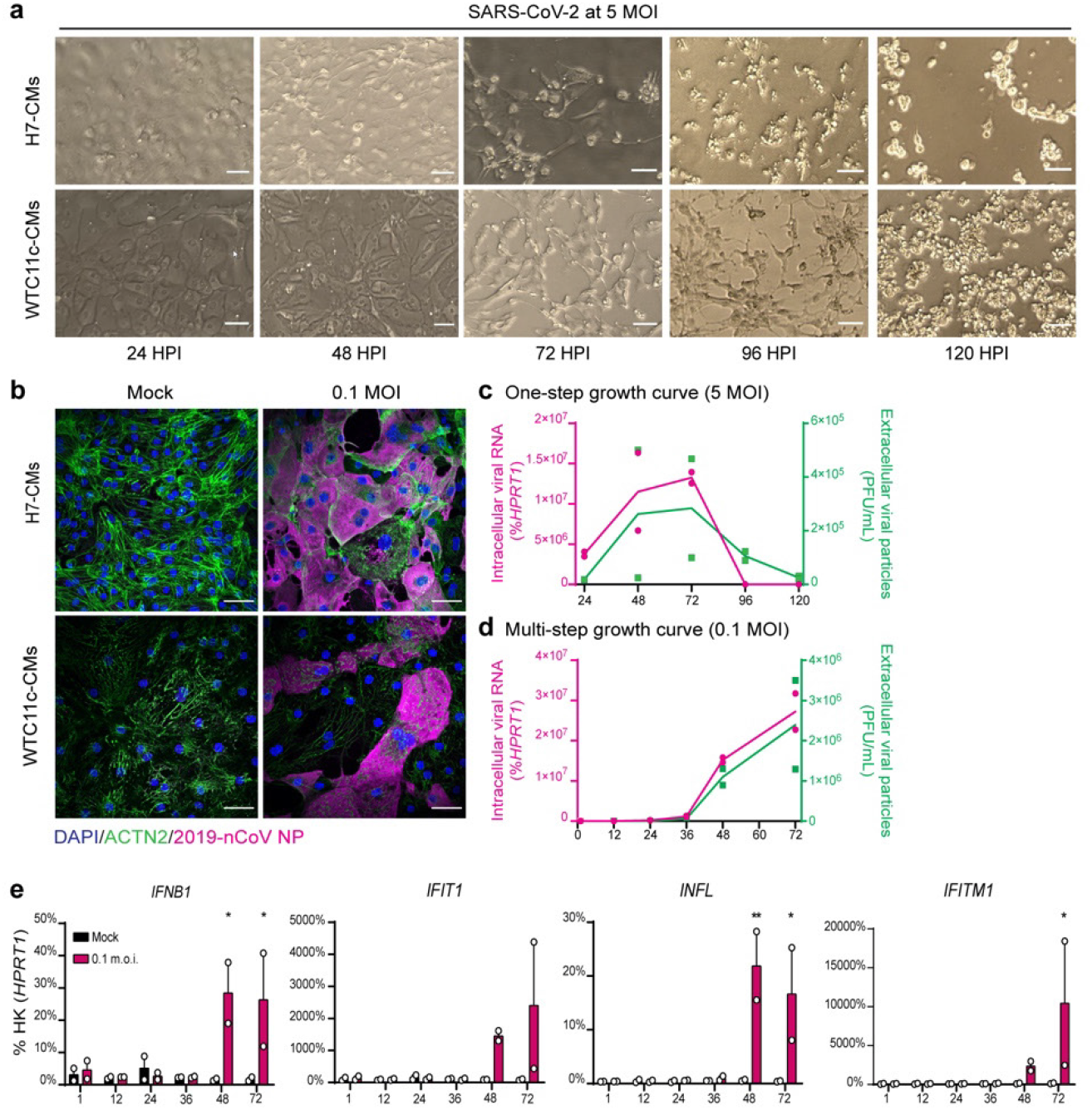
hPSC-CMs are permissive to SARS-CoV-2 infection and replication. (**a**) Cytopathic effects of SARS-CoV-2 at 5 MOI in H7 hESC-CMs and WTC11c hiPSC-CMs during a time course of 120 h. Scale bars: 100 μm. (**b**) Immunofluorescent staining of H7 hESC-CMs and WTC11c hiPSC-CMs at 48 HPI with SARS-CoV-2 dose of 0.1 MOI. Scale bars: 50 μm. Individual channels are shown in Supplementary Fig. 2c. (**c**) One-step viral growth curve in H7 hESC-CMs infected with SARS-CoV-2 at 5 MOI over a time course of 120 h. (**d**) Multi-step viral growth curve in H7 hESC-CMs infected with SARS-CoV-2 at 0.1 MOI over a time course of 72 h. For both c and d, lines connect the means of two independent experiments. Viral RNA indicating intracellular viral replication is plotted on the left y axis as % of *HPRT1*. Viral particles secreted in the supernatant are plotted on the right y axis as PFU/mL. (**e**) RT-qPCR of interferon response genes in H7 hESC-CMs infected with SARS-CoV-2 at 0.1 MOI. Mean ± SEM of 2 independent experiments. Differences *versus* non infected (mock) control by two-way ANOVA followed by Sidak correction for multiple comparisons (* = p <0.05; ** = p < 0.01).

To investigate if hPSC-CMs are permissive to SARS-CoV-2 replication, we quantified extracellular viral particles and intracellular viral RNA (by plaque assay and RT-qPCR, respectively). The one-step growth curve after 5 MOI infection indicated that significant viral replication occurred steadily from 24-72 HPI, followed by a precipitous decline as the cells died (Fig. 2c). With a multi-step growth curve in cells (0.1 MOI infection), we confirmed that SARS-CoV-2 replicated inside hPSC-CMs (Fig. 2d), with a marked increase in viral particles and RNA at 48 and 72 HPI (at which point the experiment was stopped). In agreement with morphological observations, H7 cardiomyocytes were more permissive to SARS-CoV-2 replication than WTC11c (Figs. 2c-d and Supplementary Figs. 2d-e), perhaps as a reflection of genetic or epigenetic differences that deserve further study.

Upon viral infection, pathogen-associated molecular patterns (PAMPs) initiate the early immune response *via* host pattern recognition receptors (PRRs). Once viruses uncoat, RIG-I-like receptors (RLRs) bind to the uncapped and double stranded viral RNA in the cytosol and trigger innate immune activation, leading to the production of type I and type III interferons and the interferon-induced antiviral response^28^. We found an increase in interferon transcripts (*INFB1* and *IFNL*) at 48 and 72 HPI in both cell lines, with a stronger effect in the more sensitive H7 cardiomyocytes (Fig. 2e and Supplementary Fig. 2f). The interferon-stimulated genes *IFIT1* and *IFITM1* were also upregulated at the latest time points. These results indicate that SARS-CoV-2 induces innate immune activation and interferon response also in hPSC-CMs^29^.

We next investigated whether SARS-CoV-2 infection impairs the function of hPSC-CMs. First, we evaluated electrophysiological properties of infected H7 hESC-CMs and WTC11c hiPSC-CMs using multi-electrode arrays (MEA) over a time course of 72 HPI at MOI of 0.1 and 5. Distinctly from our earlier experiments on hPSC-CM monolayers, in this context we did not observe obvious cell death in any of the conditions (Fig. 3a and Supplementary Fig. 3a). This outcome may reflect a decreased efficiency of viral infection as hPSC-CMs are plated at high density to ensure robust assessment of physiologically relevant electrophysiological properties. Remarkably, even in the absence of mayor cytopathic effects, SARS-CoV-2 infection rapidly resulted in reduced beating rate, lower depolarization spike amplitude, and decreased electrical conduction velocity (Figs. 3b-c and Supplementary Figs. 3b-c). In H7 hESC-CMs we also observed a time-dependent increase in the duration of the field potential (FPD) both in spontaneously beating and electrically paced cultures (Fig. 3d; similar measurements could not be reliably obtained from WTC11c hiPSC-CMs due to the limited amplitude of the repolarization wave after SARS-CoV-2 infection, Supplementary Fig. 3b). The FPD reflects the interval between membrane depolarization and repolarization, and as such represent an *in vitro* surrogate of the QT interval measured by an electrocardiogram. It is well known that prolongation of the QT interval is pro-arrhythmogenic^30^. Overall, abnormalities in the generation and propagation of electrical signals were significant even in the absence of substantial cell death, suggesting that SARS-CoV-2 infection in cardiomyocytes could directly create a substrate for arrhythmias. These properties of SARS-CoV-2 infection in cardiomyocytes could explain the high rate of arrhythmia (~14%) which has been observed in COVID-19 patients^5^.

**Figure 3.**
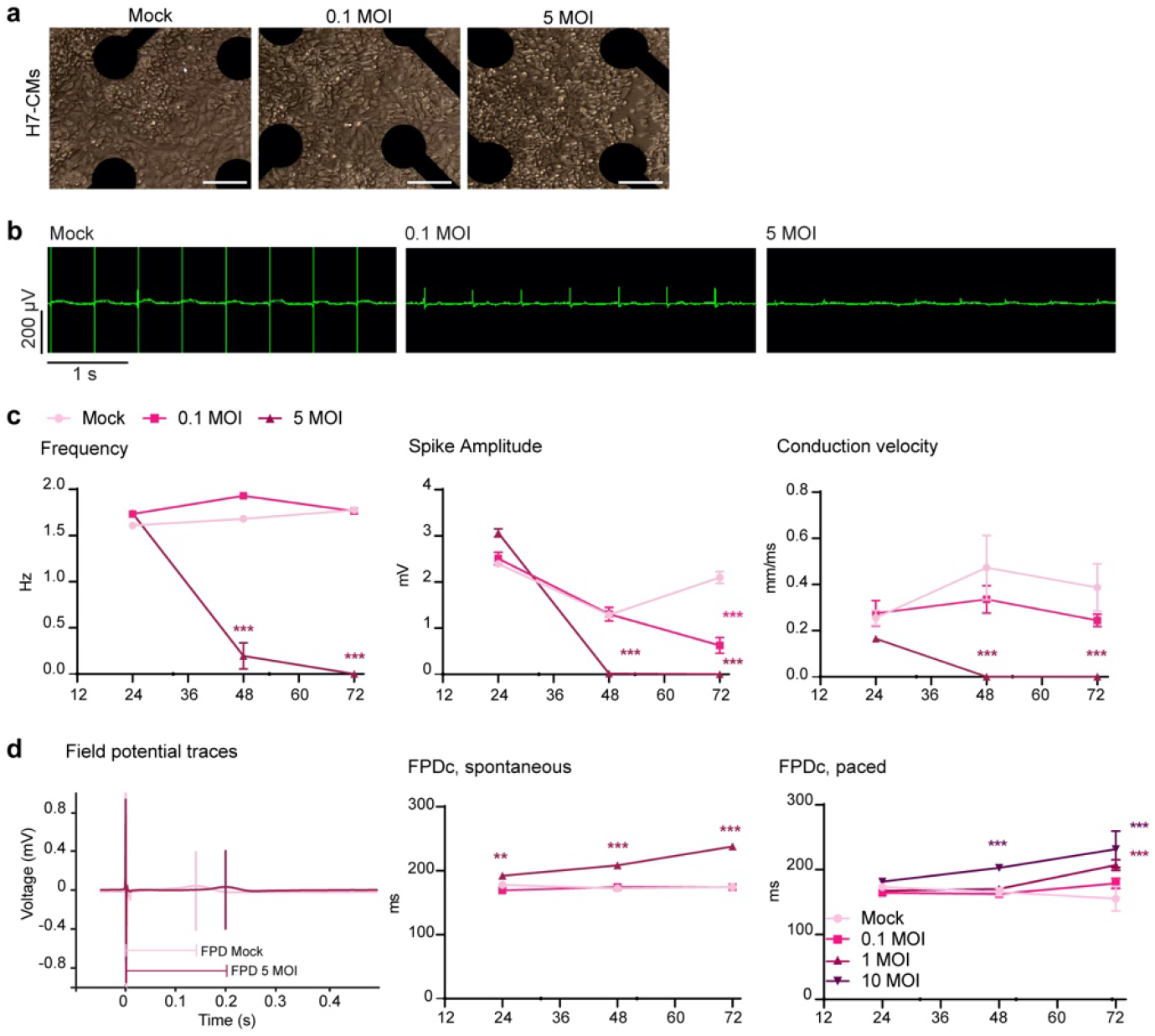
Electrophysiological alterations in hPSC-CMs infected with SARS-CoV-2. (**a**) Representative images of SARS-CoV-2-infected H7 hESC-CMs on MEA wells at 72 HPI. Scale bars: 50μm (**b**) Representative recordings of spontaneous electrical activity of SARS-CoV-2-infected H7 hESC-CMs at 72 HPI. (**c**) Representative quantifications of electrophysiological properties from MEA analyses in SARS-CoV-2-infected H7 hESC-CMs. Mean ± SEM of 8 wells. Differences *versus* mock control by two-way ANOVA with Sidak correction for multiple comparisons (*** = p < 0.001). (**d**) Representative field potential traces in SARS-CoV-2-infected H7 hESC-CMs at 72 HPI, and quantification of field potential duration corrected by beat rate (FPDc) in spontaneous and paced experiments. Mean ± SEM of 8 and 6 wells for spontaneously beating and paced cells, respectively. Statistical analyses as for panel c (** = p < 0.01).

We then evaluated the contractile properties of hPSC-CMs using three-dimensional engineered heart tissues (3D-EHTs), following their contractile behavior through magnetic field sensing^31^ (Figs. 4a-b). For these experiments we focused on WTC11c hiPSCs since 3D-EHTs from H7 hESC-CMs proved to spontaneously beat at too high a frequency (> 2 Hz) to enable accurate measurements of contractile behavior (i.e. the tissue had a tetanic-like contraction with minimal relaxation between beats at this frequency). We infected 3D-EHTs from WTC11c hiPSC-CMs with 10 MOI (to facilitate infection within the non-vascularized, cell-dense tissue), and analyzed their contraction for a week. The maximal twitch force decreased as early as 72 HPI, and the contractions continued to subside to < 25% of original force at 144 HPI (Figs. 4c-d, and Supplementary Videos 1 and 2). Overall, the significant impairment in the contractile properties of 3D-EHTs demonstrates that the mechanical function of cardiomyocytes is impacted by SARS-CoV-2 infection, and this could contribute to whole-organ cardiac dysfunction in patients^32^.

**Figure 4.**
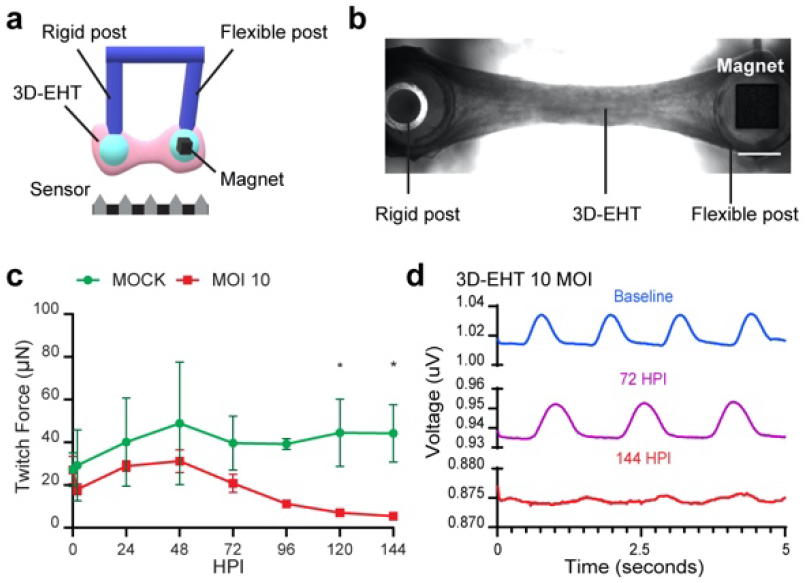
Force production of 3D-EHTs made from hiPSC-CMs progressively declines during SARS-CoV-2 infection. (**a**) Schematic representation of the magnetic sensing system: the 3D-EHT is suspended between two posts (one rigid, one flexible). The magnet is localized inside the flexible post, and the post movement during 3D-EHT contraction is recorded by the sensor located at the bottom of the dish (converting displacement into voltage changes). (**b**) Representative picture of a 3D-EHT. Scale bar: 1 mm. (**c**) Representative time course analysis of twitch force in 3D-EHTs from WTC11c hiPSC-CMs after SARS-CoV-2 infection at 10 MOI. Data are shown as mean ± SEM for mock controls and 4 infected 3D-EHTs. Differences *versus* mock by two-way ANOVA with Sidak correction for multiple comparisons (* = p < 0.05). (**d**) Representative voltage traces at different time points.

A rapidly increasing number of reports acknowledge cardiovascular involvement as a prevalent complication observed in COVID-19 patients, but discriminating between direct *versus* indirect effects is still an open challenge^12,14,18,19^. Our results provide important evidence that SARS-CoV-2 has the ability to directly infect cardiomyocytes, to impair both their electrophysiological and contractile properties, and eventually to induce cell death. These results support the hypothesis that, independent of inflammation or coagulopathy, SARS-CoV-2 can cause direct functional heart damage.

One limitation of this study is our reliance on hPSC-CMs, which are well-known for their functional immaturity^33^. Moreover, while the *in vitro* systems we used have been successfully leveraged to model electrophysiological and contractile alterations due to drugs or inherited mutations^34,35^, their application to modeling COVID-19 will require further validation. A recent report by Dolhnikoff et al. identified coronaviral particles in the cytoplasm of cardiomyocytes, endothelial cells, and fibroblasts by electron microscopy in the heart of an 11 year-old child who died from multi-system inflammatory syndrome in children (MIS-C) following COVID-19 infection^20^. This indicates that *in vivo* cardiomyocytes with substantially greater maturity than used here are susceptible to SARS-CoV-2 infection. COVID-19 patients are commonly treated with steroids to control systemic inflammation. However, our data suggests that treatments aimed to control the direct damage of SARS-CoV-2, e.g. antiviral medications and/or cardioactive drugs, should also be taken into consideration to prevent long-term cardiovascular complications.

## Methods

### Cell culture

Undifferentiated RUES2 hESCs (RUESe002-A; WiCell) and WTC11c hiPSCs (a gift of Dr. Bruce Conklin, Gladstone Institutes, San Francisco) were maintained in mTeSR1 (Stemcell Technologies) on tissue culture dishes coated with Matrigel (Corning) at 0.17 mg/mL, and passaged as small clumps using Versene (Gibco). Cardiomyocytes from RUES2 and WTC11c were differentiated as previously described^36^. Briefly, undifferentiated cells were seeded at 1,000 cells/cm^2^ in mTeSR1 supplemented with 10 μM Y-27632 (Tocris) on Matrigel-coated dishes. After 24 h media was changed to mTeSR1 with 1 μM CHIR-99021 (Cayman). On day 0 mesoderm differentiation was induced with 3 μM CHIR-99021 in RPMI-1640 media (ThermoFisher) supplemented with 500μg/mL BSA (Sigma-Aldrich) and 213μg/mL ascorbic acid (Sigma-Aldrich), denoted as RBA media. On day 2 media was changed to RBA containing 2 μM WNTC59 (Selleckchem). On day 4 media was changed to plain RBA. On day 6 media was changed to RPMI-1640 plus B-27 supplement (ThermoFisher), with further media changes every other day. Heat-shock was performed on day 13 for 30 min at 42 °C, and on day 14 cardiomyocytes were dissociated and frozen in CS10 cryopreservation media (Sigma-Aldrich). H7 hESCs (WA07; WiCell) were differentiated in suspension culture format by collaborators at the Center for Applied Technology Development at the City of Hope in California (as previously described^37^), received by dry-ice shipment, and stored in liquid nitrogen before use.

### Flow cytometry

Cardiac differentiation efficiency was determined by flow cytometry in hPSC-CMs fixed with 4% paraformaldehyde for 15 min at room temperature. Cells were centrifuged at 300 g for 5 min and incubated at room temperature for 1 h with either APC-cTnT antibody (Miltenyi Biotech #130-120-543) or APC-IgG1 isotype control antibody (Miltenyi Biotech #130-120-709), both used at 1:100 dilution in DPBS (Gibco) with 5% fetal bovine serum (FBS)and 0.75% saponin. Following washes with DPBS with 5% FBS, samples were run on a BD FACSCanto II flow cytometer and data from 10,000 valid events were acquired with the BD FACSDIVA software. Analysis was performed with FlowJo v10.7.

### RNA-seq

Bulk RNA-seq datasets from differentiating RUES2 hESC-CMs had been previously generated and analyzed^36^ (GEO dataset: GSE106688).

### Western blot

Cell pellets were incubated in RIPA Buffer supplemented with 1X Protease inhibitors (ThermoFisher) at 4 °C for 20 min. Samples were then centrifuged at 21,000 g for 15 min at 4 °C, and protein concentration in the supernatant was quantified with BCAssay (ThermoFisher). 25 μg of protein were mixed with 1X non-reducing SDS sample buffer and incubated at 37 °C for 30 min. Samples were run on 4-20% mini-PROTEAN TGX precast gels (Bio-Rad) and then transferred on PVDF membranes. Membranes were incubated with 5% nonfat dry milk in TBS buffer supplemented with 0.1% Tween-20 (blocking buffer) for 1 h at room temperature. Primary antibodies were incubated in blocking buffer for 2 h at room temperature (rabbit anti-ACE2 [Abcam #ab15348, used at 1:500 dilution]; mouse anti-GAPDH [Abcam #ab8245, used at 1:3,000 dilution]). Membranes were washed and further incubated with fluorescent dye-conjugated secondary antibodies for 1 h at room temperature (AlexaFluor 647 goat anti-rabbit IgG1 and AlexaFluor 488 goat anti-mouse IgG1, both used at 1:1,000 dilution in blocking buffer) and fluorescent signals were acquired using with a GelDoc Imager (Bio-Rad).

### Single cell RNA-seq

A single cell suspension was generated from cultures of RUES2 hESC-CMs at day 30 of differentiation, and single cell RNA-seq was performed using the Chromium NextGEM Single Cell 3’ kit (10X Genomics). 10,000 cells per condition were loaded on independent microfluidics channels to generate Gel bead-in-Emulsion (GEMs), which were further processed as per the manufacturer’s instructions to generate Illumina-compatible sequencing libraries. The sample was analyzed using two runs of high output NextSeq 500 with a 150 cycle kit, reading 28 base pairs for read 1 (barcode and UMI), 91 base pairs for read 2 (3’ end of cDNAs), and 7 base pairs for the i7 index. For data analysis, cellranger mkfastq was used to transfer demultiplexed raw base call files into library-specific FASTQ files. The FASTQ files were separately mapped to the GRC38 human reference genome using STAR as a part of the cellranger pipeline. Gene expression counts were done using cellranger count based on Gencode v25 annotation, and cell identifiers and Unique Molecular Identifiers (UMI) were filtered and corrected with default setting. Raw cellranger count outputs was aggregated and visualized by a subsampling procedure using cellranger aggr. Downstream analysis was performed in the R package Seurat^38,39^. Filters were applied to eliminate cells with less than 1,000 genes detected, with over 40,000 UMIs, or with over 35% mitochondrial gene reads. Post filtering, cell-to-cell gene expression was normalized by total expression, multiplied by the scale factor of 10000, and the result is log-transformed [Log norm exp]). Data dimensionality reduction was done by principal component analysis on the top 2,000 most variable genes. The top 10 principal components (PCs) that explained most variance were selected, as confirmed by an Elbow plot. For visualization, UMAP dimensionality reduction for the top 10 PCs was performed to produce coordinates for cells in 2 dimensional space. The relative expression levels of genes of interest were plotted using the FeaturePlot function in Seurat.

### SARS-CoV-2 generation

All experiments using live virus were performed in the Biosafety Level 3 (BSL-3) facility at the University of Washington in compliance with the BSL-3 laboratory safety protocols (CDC BMBL 5^th^ ed.) and the recent CDC guidelines for handling SARS-CoV-2. Before removing samples from BSL-3 containment, samples were inactivated by Trizol or 4% paraformaldehyde, and the absence of viable SARS-CoV-2 was confirmed for each sample by plaque assays. SARS-Related Coronavirus 2, Isolate USA-WA1/2020 (SARS-CoV-2) was obtained from BEI Resources (NR-52281) and propagated in VERO cells (USAMRIID). Briefly, VERO cells were maintained in DMEM (Gibco) supplemented with 10% heat-inactivated FBS, 100 U/mL penicillin, and 100 U/mL streptomycin at 37 °C in a 5% CO_2_ humidified incubator. To generate virus stock, cells were washed once with DPBS and infected with SARS-CoV-2 in serum-free DMEM. After 1 h of virus adsorption, the inoculum was replaced with DMEM supplemented with 2% heat-inactivated FBS, and cells were incubated at 37 °C in a 5% CO_2_ incubator until ~70% of cells manifested cytopathic effects. The virus was harvested by collecting the culture supernatant followed by centrifugation at 3,000 g for 15 min at 4 °C to remove the cell debris. Virus titer was then measured by plaque assay on VERO cells (as described below), and stocks were stored at −80°C.

### SARS-CoV-2 titering

Viral preparations and culture supernatant from SARS-CoV-2-infected cardiomyocytes were titered using a plaque assay. Briefly, 350,000 VERO cells were seeded in 12-well plates and incubated for 1 h at 37 °C with 10-fold dilutions of virus-containing media. A solution of 1:1 1% agarose and 2X DMEM supplemented with 4% heat-inactivated FBS, L-glutamine, 1X antibiotic-antimycotic (Gibco), and 220 mg/mL sodium pyruvate was layered on top of the cells, followed by incubation at 37 °C for 2 days. After fixing with 10% formaldehyde, the agarose layer was removed and cells were stained with 0.5% crystal violet solution in 20% ethanol. Plaques were counted, and the virus titer in the original sample was assessed as plaque-formation unit per mL (PFU/mL).

### Viral infection

Cryopreserved hPSC-CMs were thawed and plated in RPMI-1640 supplemented with B-27, 5% FBS, and 10 μM Y-27632. After 24 h the media was replaced with RPMI-1640 supplemented with B-27 only. After 3 days, cardiomyocytes were harvested with Versene supplemented with 0.5% Trypsin (Gibco) at 37 °C for 5 min to obtain single-cell suspensions. 250,000 cardiomyocytes were seeded in Matrigel-coated 12-well plates in RPMI-1640 supplemented with B-27, 5% FBS, and 10 μM Y-27632. The media was replaced with RPMI-1640 supplemented with B-27 the next day, and then every other day for 1 week. SARS-CoV-2 was diluted to the desired MOI in DMEM and incubated on hPSC-CMs for 1 h at 37 °C (non-infected [mock] controls were incubated with DMEM only). Cells were then washed with DPBS and cultured in RPMI-1640 supplemented with B-27.

### Immunofluorescence

200,000 cardiomyocytes were plated on glass-bottom 24-well plate (CellVis) and infected as described above. Cells were fixed with 4% paraformaldehyde in DPBS for 30 min at room temperature and then washed 3 times with DPBS for 5 min. Cells were permeabilized using 0.25% Triton X-100 (Sigma-Aldrich) in DPBS and blocked for 1 h with 10% normal goat serum supplemented with 0.1% Tween-20 in DPBS. Primary antibodies were incubated overnight at 4 °C in DPBS with 1% normal goat serum and 0.1% Tween-20 (rabbit anti-2019-nCoV NP [Sino Biological #40143-R019, used at 1:200 dilution]; and mouse anti-Sarcomeric α-actinin [Abcam ab# ab9465, used at 1:500 dilution]). Cell were washed three times with DPBS containing 0.2% Tween-20, and incubated for 1 h at room temperature with secondary antibodies diluted in DPBS supplemented with 1% BSA and 0.1% Tween-20 (AlexaFluor 594 goat anti-rabbit IgG1 and AlexaFluor 647 goat anti-mouse IgG1, both used at 1:1,000 dilution). DAPI (Sigma-Aldrich) was diluted at 300 nM in water and incubated on the cells for 15 min at room temperature, followed by three washes in DPBS containing 0.2% Tween-20. Images were taken with a 40x oil objective on a Nikon Eclipse microscope with Yokogawa W1 spinning disk head, and formatted with Fiji software.

### Gene expression analysis and viral RNA detection

Infected cardiomyocytes were washed once with DPBS and incubated with 400 μL per well of Trizol reagent (Invitrogen) for 10 min at room temperature. Chloroform was added in a 5:1 ratio to Trizol, and samples were incubated at room temperature for 2 min. The aqueous phase was separated by centrifugation (21,000 g for 15 min at 4 °C) and incubated with isopropanol (1:1 ratio) and 25 μg of Glycoblue (ThermoFisher) for 10 min at room temperature. RNA pellets were harvested by centrifugation (21,000 g for 15 min at 4 °C), washed twice with 75% ethanol, and resuspended in nuclease-free water. cDNA was prepared with M-MLV reverse transcriptase according to the manufacturers’ instruction. Quantitative real-time reverse transcription PCR (RT-qPCR) was performed with SYBR Select Master Mix (Applied Biosystems) using 10 ng of cDNA and 400 nM forward and reverse primers (Supplementary table 1). Reactions were run on a CFX384 Real-Time System (Bio-Rad), and data was analyzed using the ΔΔCt method using *HPRT1* as the housekeeping gene. Primers were designed using PrimerBlast, and confirmed to amplify a single product.

### Electrophysiological analysis with MEA

Cryopreserved cardiomyocytes were thawed and cultured as described above. CytoView MEA 48- and 24-well plates (Axion BioSystems) were coated with 0.17 mg/mL of Matrigel for 1 h at 37 °C. 50,000 (48-well plate) or 100,000 (24-well plate) hPSC-CMs were resuspended in 6 μL or 10 μL, respectively, and plated on each MEA well, as previously described^40^. Media was changed with RPMI-1640 supplemented with B-27 every other day for 1 week. One the day of viral infection, cells were washed once with DPBS and incubated with 50 μL (48-well plate) or 100 μL (24-well plate) of SARS-CoV-2 suspension for 1 h at 37 °C. Media was replaced with RPMI-1640 supplemented with B-27, and the plate was transferred directly into Maestro Pro system (Axion BioSystems) and kept at 37 °C with 5% CO_2_ for the duration of the experiment. Electrophysiological recordings were taken for 5 min at specified time points using Axis software version 2.0.4. (Axion BioSystems). Voltage was acquired simultaneously for all the electrodes at 12.5 kHz, with a low-pass digital filter of 2 kHz for noise reduction. The beat detection threshold was 100 μV, and the FPD detection used a polynomial regression algorithm with the threshold set at 1.5 × noise to detect repolarization waves. Pacing was performed at 2 Hz with an alternating square wave (± 1 V, 100 nA, 8.33 kHz) delivered through the dedicated stimulator in the Maestro Pro system to a selected electrode (not used for recording). Automated analysis was performed using Cardiac Analysis Software v3.1.8 (Axion BioSystems), which automatically computes the Fridericia correction to account for beat rate variability during FPD measurements [FPDc = FPD/(beat period)^1/3^].

### Contractility analysis with 3D-EHTs

3D-EHTs were generated from hPSC-CMs embedded with stromal cells in a 3D fibrin gel suspended between pairs of silicone posts, as previously described^31^. For each pair of silicone posts one was flexible and had a 1 mm^3^ magnet embedded in its tip, and the other post was rendered rigid by embedding a 1.1 mm glass capillary tube. Each 3D-EHT was casted in a mold made of 2% agarose by adding 500,000 WTC11c hiPSC-CMs and 50,000 HS27a stromal cells in a fibrin gel solution (89 μL RPMI-1640 supplemented with B-27, 5.5 μl of DMEM/F12, [Gibco], 2.5 μL of 200 mg/mL bovine fibrinogen [Sigma-Aldrich], and 3 μL of 100 U/mL thrombin [Sigma-Aldrich]. The cell-gel mixture was incubated at 37 °C for 2 h to allow for fibrin polymerization. Afterwards, 3D-EHTs were transferred from the agarose molds to 24-well tissue culture dishes containing 3D-EHT media (RPMI-1640 media supplemented with B-27 and 5 mg/mL aminocaproic acid [Sigma-Aldrich]. Media was changed every other day for 2 weeks. For SARS-CoV-2 infection, 3D-EHTs were temporarily housed in 2% agarose molds, and 200 μL of viral solution in DMEM was used to infect each single tissue for 1 h at 37 °C (DMEM was used for mock controls). 3D-EHTs were then transferred in fresh 3D-EHT media for the rest of the experiment. Twitch force was recorded by tracking the movement of magnets embedded in the flexible posts, as previously described^31^. Briefly, we used a custom-built printed circuit board (PCB) containing giant magnetoresistive (GMR) sensors (NVE, Eden Prairie, MN) in a Wheatstone bridge configuration and relying on instrumentation amplifiers and operational amplifiers to filter out signal noise. 3D-EHTs in the 24-well dish were placed into a 3D-printed caddy that contained the PCB with GMR sensors such that the flexible, magnetic posts of 3D-EHTs were directly above each GMR sensor. Data from the PCB was collected by LabView (National Instruments) on a laptop in the BSL-3 facility. The voltage traces from the magnetic sensors were analyzed for amplitude and frequency using a custom Matlab protocol. The amplitudes were then converted from voltage to twitch force using a characterization constant.

### Statistical analyses

Statistical analyses were performed using Prism 8.1.3 (GraphPad). The type and number of replicates, the statistics plotted, the statistical test used, and the test results are described in the figure legends.

## Acknowledgements

We thank Drs. Farid Moussavi-Harami, Kenta Nakamura, and Daniel Yang for helpful discussions during the development of this project, and Dr. Aidan Fenix for experimental support. We are grateful to Axion Biosystem for proving material and assistance for the MEA experiments. This research was assisted by the University of Washington Cell Analysis Facility (Department of Immunology), and by the Lynn and Mike Garvey Cell Imaging Core (Institute for Stem Cell and Regenerative Medicine). We would also like to acknowledge the University of Washington BSL-3 facility, which provided assistance during experiments and ensured our safety. SM is supported by a Postdoctoral Fellowship from the UW Institute for Stem Cell and Regenerative Medicine. This work was supported in part by the National Science Foundation (CMMI-166173 to N.J.S.), National Institutes of Health (R01 HL149734 to N.J.S.; R01 AI118916, R01 AI127463, R01 AI145296, R21 AI145359, U01 AI151698, UM1 AI148684, and U19 AI100625 to M.G.; R01 HL128362 and R01 HL146868 to C.E.M.), and from State of Washington and philanthropical support to the UW Institute for Stem Cell and Regenerative Medicine.

## Author contributions

S.M. performed hiPSC-CM differentiation, viral infections, western blots, viral plaque, MEA and 3D-EHTs assays, and wrote the first draft of the manuscript. T.Y.H. cultured and expanded SARS-CoV-2 and performed immunofluorescence. T.H. designed and fabricated the 3D-EHTs magnetic sensing system, and analyzed the data. A.K. performed and analyzed RT-qPCR. H.R. contributed to experimental design and differentiated hiPSC-CMs. X.Y. contributed to experimental design and analyzed sc-RNA-seq data. L.P. contributed to experimental design and supervised the experiments. A.B. contributed to the study’s conception, experimental design and execution, and assisted in writing the manuscript. N.J.S., M.G. and C.E.M. conceived and supervised the study, obtained research funding and contributed to data analysis and writing the manuscript.

