## Supplementary information for "SARS-CoV-2 infects human pluripotent stem cell-derived cardiomyocytes, impairing electrical and mechanical function"

**Supplementary Figure 1 – hPSC-CMs express SARS-CoV-2 receptors and processing factors.**

**Supplementary Figure 2 – hPSC-CMs are permissive to SARS-CoV-2 infection and replication.**

**Supplementary Figure 3 – Electrophysiological alterations in hPSC-CMs infected with SARS-CoV-2.**

**Supplementary Table 1 – RT-qPCR primer sequences.**

**Supplementary Video 1 – 3D-EHT before SARS-CoV-2 infection (online).**

**Supplementary Video 2 – SARS-CoV-2 infected 3D-EHT at 144 HPI (online).**

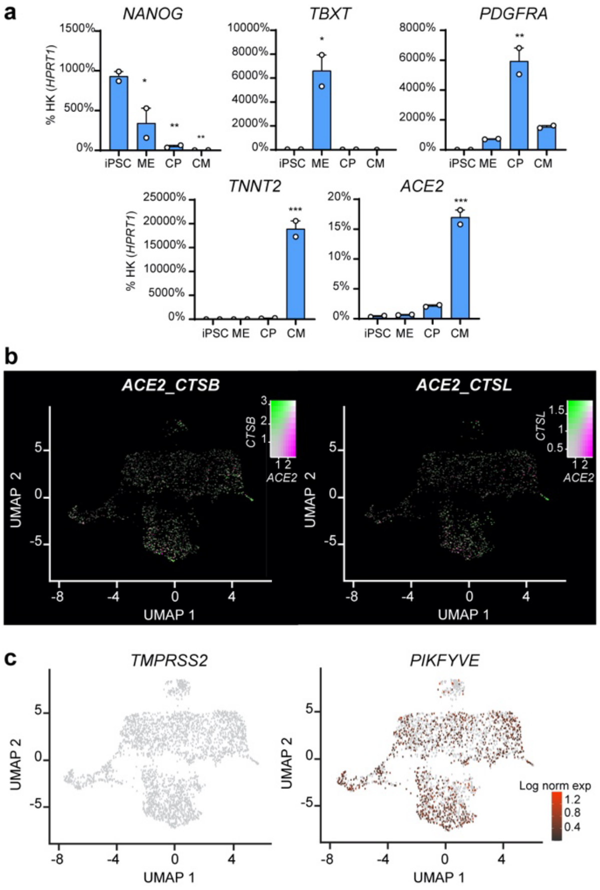


**Supplementary Figure 1 – hPSC-CMs express SARS-CoV-2 receptors and processing factors.** (**a**) RT-qPCR analysis during WTC11c hiPSC-CM differentiation. iPSC: induced pluripotent stem cells (day 0); ME: mesoderm (day 2); CP: cardiac progenitors (day 5); CM: cardiomyocytes (day 14). Differences versus iPSC by one-way ANOVA with Sidak correction for multiple comparisons (* = $p < 0.05$; ** = $p < 0.01$; *** = $p < 0.001$). (**b-c**) sc-RNA-seq gene expression heatmaps from RUES2 hESC-CMs after dimensionality reduction through UMAP. In b, plots showcase double positive cells for ACE2 and CTSL, and ACE2 and CTSB.

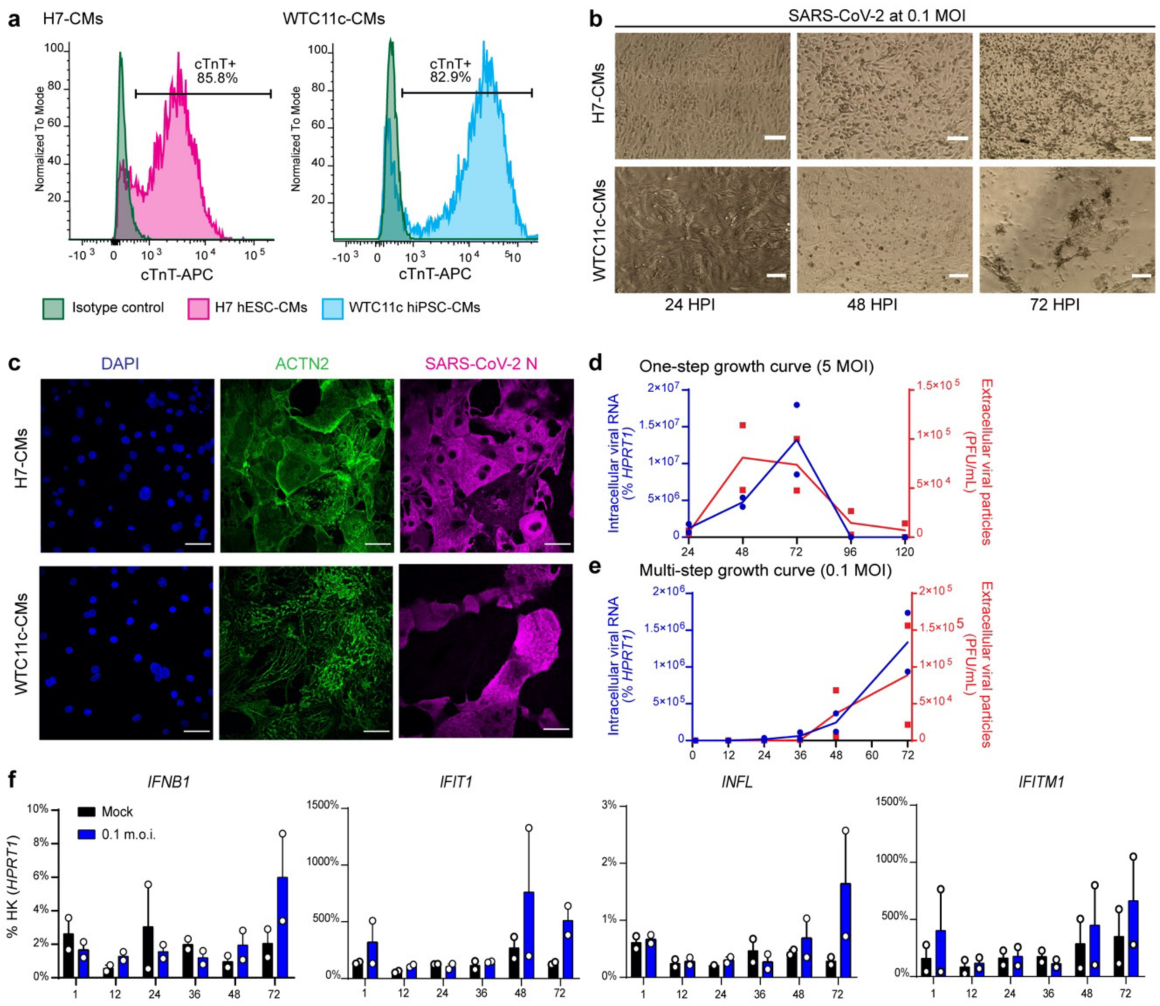


**Supplementary Figure 2 – hPSC-CMs are permissive to SARS-CoV-2 infection and replication.** (**a**) Flow cytometry analysis for cardiac troponin T (cTnT) expression in H7 hESC-CMs and WTC11c hiPSC-CMs. (**b**) Representative images of H7 hESC-CMs and WTC11c hiPSC-CMs infected with SARS-CoV-2 at 0.1 MOI during a time course of 72 h. Scale bars: 100 µm. (**c**) Single channel images for the immunostainings of SARS-CoV-2-infected hPSC-CMs shown in Figure 2b. (**d**) One-step viral growth curve in WTC11c hiPSC-CMs infected with SARS-CoV-2 at 5 MOI. (**e**) Multi-step viral growth curve in WTC11c hiPSC-CMs infected with SARS-CoV-2 at 0.1 MOI. For both e and d, lines connect the mean of two independent experiments. Viral RNA indicating intracellular viral replication is plotted on the left y axis as % of HPRT1. Viral particles secreted in the supernatant are plotted on the right y axis as PFU/mL. (**f**) RT-qPCR of interferon response genes in WTC11c hiPSC-CMs infected with SARS-CoV-2 at 0.1 MOI. Mean ± SEM of 2 independent experiments.

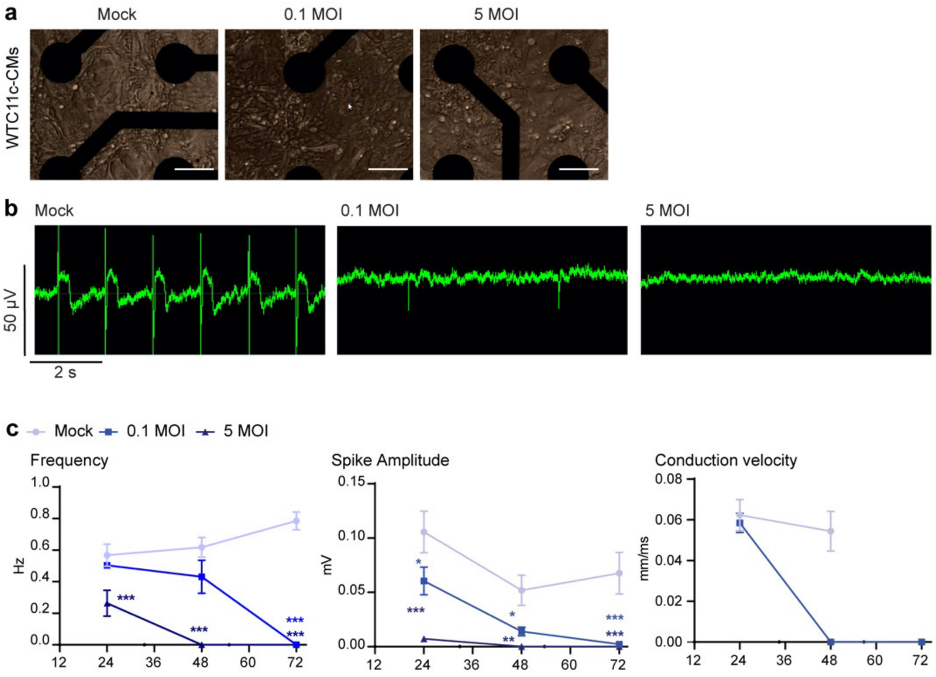


**Supplementary Figure 3 – Electrophysiological alterations in hPSC-CMs infected with SARS-CoV-2.** (**a**) Representative images of SARS-CoV-2-infected WTC11c hiPSC-CMs on MEA well at 72 HPI. Scale bars: 50 µm. (**b**) Representative recording of spontaneous electrical activity of SARS-CoV-2-infected WTC11c hiPSC-CMs at 72 HPI. (**c**) Representative quantification of electrophysiological properties from MEA analyses on SARS-CoV-2-infected WTC11c hiPSC-CMs. Mean ± SEM of 8 wells. Differences versus mock calculated by two-way ANOVA with Sidak correction for multiple comparisons (* = $p < 0.05$; ** = $p < 0.01$; *** = $p < 0.001$).

**Supplementary Table 1 – RT-qPCR primer sequences**

| Target | Forward primer (5' – 3') | Reverse primer (5' – 3') |
|---|---|---|
| *HPRT1* | TGACACTGGCAAAACAATGCA | GGTCCTTTTCACCAGCAAGCT |
| *NANOG* | TTTGTGGGCCTGAAGAAAACT | AGGGCTGTCCTGAATAAGCAG |
| *TBXT* | CAAATCCTCATCCTCAGTTTG | GTCAGAATAGGTTGGAGAATTG |
| *PDGFRA* | GCTCACTTCACTCTCCCCAAAG | CCGGCGTTCCTGGTCTTAG |
| *TNNT2* | TTCACCAAAGATCTGCTCCTCGCT | TTATTACTGGTGTGGAGTGGGTGTGG |
| *ACE2* | CCATCAGGATGTCCCGGAG | TGGAGGCATAAGGATTTTCTCCA |
| *SARS-CoV-2_E* | GAACCGACGACGACTACTAGC | ATTGCAGCAGTACGCACACA |
| *IFIT1* | AGAAGCAGGCAATCACAGAAAA | CTGAAACCGACCATAGTGGAAAT |
| *IFITM1* | TACTCCGTGAAGTCTAGGGACAG | AACAGGATGAATCCAATGGTCA |
| *IFNB* | CTTGGATTCCTACAAAGAAGCAGC | TCCTCCTTCTGGAACTGCTGCA |
| *IFNL* | AACTGGGAAGGGCTGCCACATT | GGAAGACAGGAGAGCTGCAACT |
